## Supplementary figures and tables for "Genetic Network Shaping Kenyon Cell Identity and Function in *Drosophila* Mushroom Bodies"

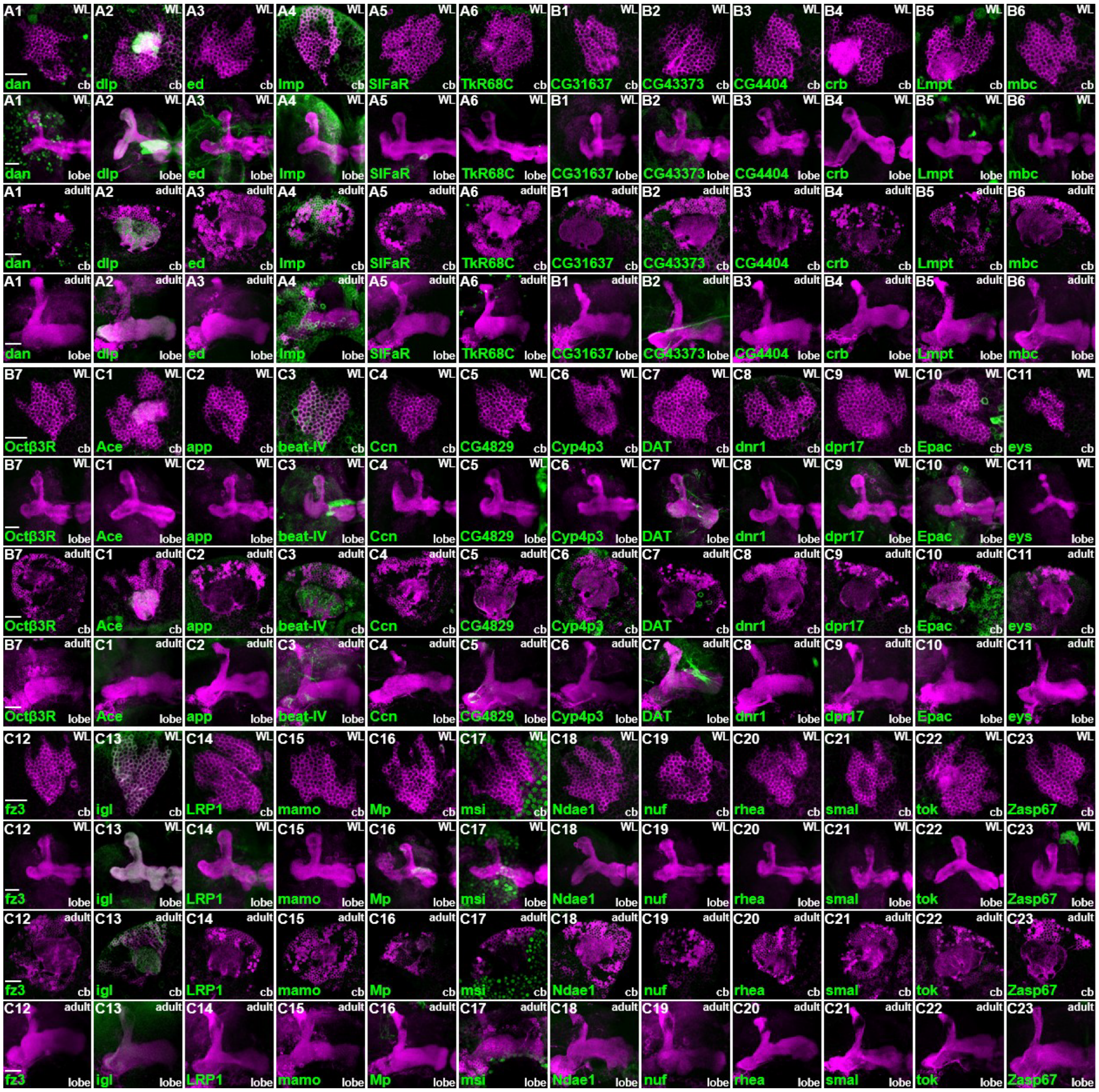


#### S1 Fig. GFP-line screen for KC subtype markers

Previous RNA-seq studies revealed genes of interest with the preferential expression in KC subtypes of adult brains [1, 2]. Among these genes, the expression levels of *ab* and *mamo* are enriched respectively in γ and α'/β' neurons [3, 4]. Therefore, these markers were utilized for the identification of genes specifically expressed in KC subtypes from RNA-seq datasets [1, 2]. To leverage the RNA-seq information to obtain freely accessible reagents for studies on KC development, KC subtype marker-expressing lines were collected, each of which carry GFP transgenes either derived from engineered BAC clones or inserted in genes of interest, from fly stock centers [5-7]. For the BAC-GFP lines, flies were generated with transgenes carrying BAC genomic DNAs of genes of interest, which contain mostly intact regulatory fragments, and an engineered DNA fragment, which permits the expression of GFP and other tags at the C-terminus of those proteins encoded by genes of interest [5]. For GFP-trapping lines, flies were generated by remobilizing or inserting transgenes for expression of GFP and tags fused in frame with proteins encoded by genes of interest [6, 7]. One strategy utilized to generate these GFP-trapping lines is by site-specifically integrating the DNA fragment which encodes in-frame GFP and tags into a coding intron of genes of interest through the Minos-mediated integration cassette (MiMIC) system [6]. To simplify this GFP-marker screen, the top list of genes enriched in γ and α/β neurons was selected using RNA-seq data from the Alyagor study to examine expression patterns [2]. Meanwhile, all genes enriched in α'/β' neurons from the RNA-seq data of the Shih study were examined for their capacity to serve as α'/β'-specific markers [1]. In total, seven, nine and twenty-four GFP-lines with specific markers were respectively identified for γ, α/β and α/'β' neurons. (A-C) In addition to the selected lines described in Fig 1, the remaining GFP-lines were depicted with the larval and adult expression patterns at cell body (cb) and lobe regions in S1 Fig. Six (panels A1-A6), seven (panels B1-B7) and twenty-three (panels C1-C23) GFP-lines were identified for potential expression in γ, α/β and α/'β' neurons, respectively (see Supplementary table 1 for the description of expression patterns). Of note, the Mamo-sfGFP-TVPTBF line (panel C15) did not appear to express in KCs. Trio was stained in magenta in panels. Scale bar: 10 μm.

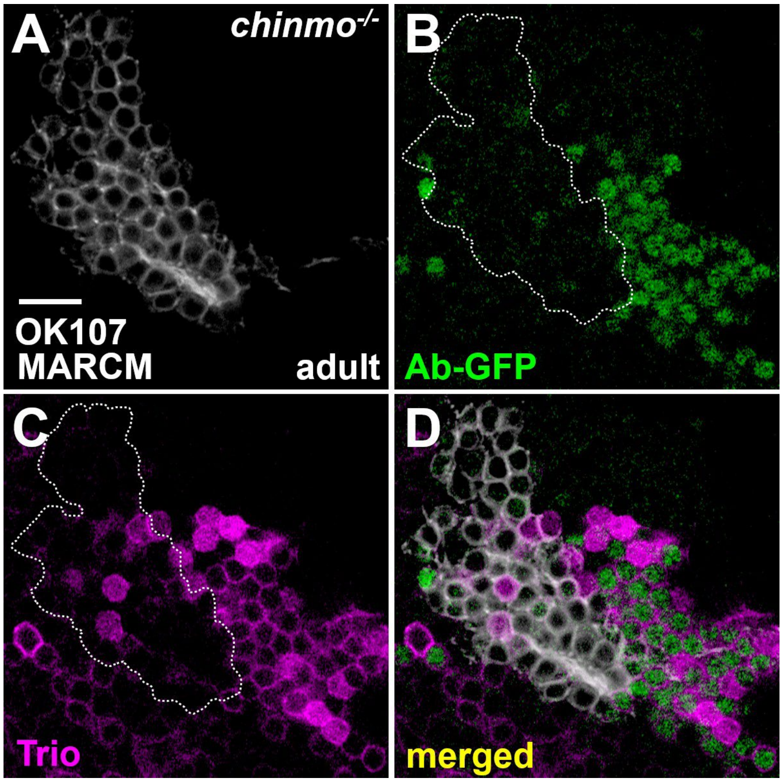


**S2 Fig. Downregulation of Ab-GFP in KCs in the *chinmo* mutation**

Ab-GFP expression (green) was compromised in KCs of *chinmo*[1] mutants in the MARCM analysis using GAL4-OK107 (white). This justifies to use Ab-GFP as the replacement of Ab antibody for the readout of a γ-specific marker. Mosaic clones were induced at newly-hatched larva and analyzed at adult brains. Trio (magenta) was used to label γ and α'/β' neurons. Scale bar: 10 μm.

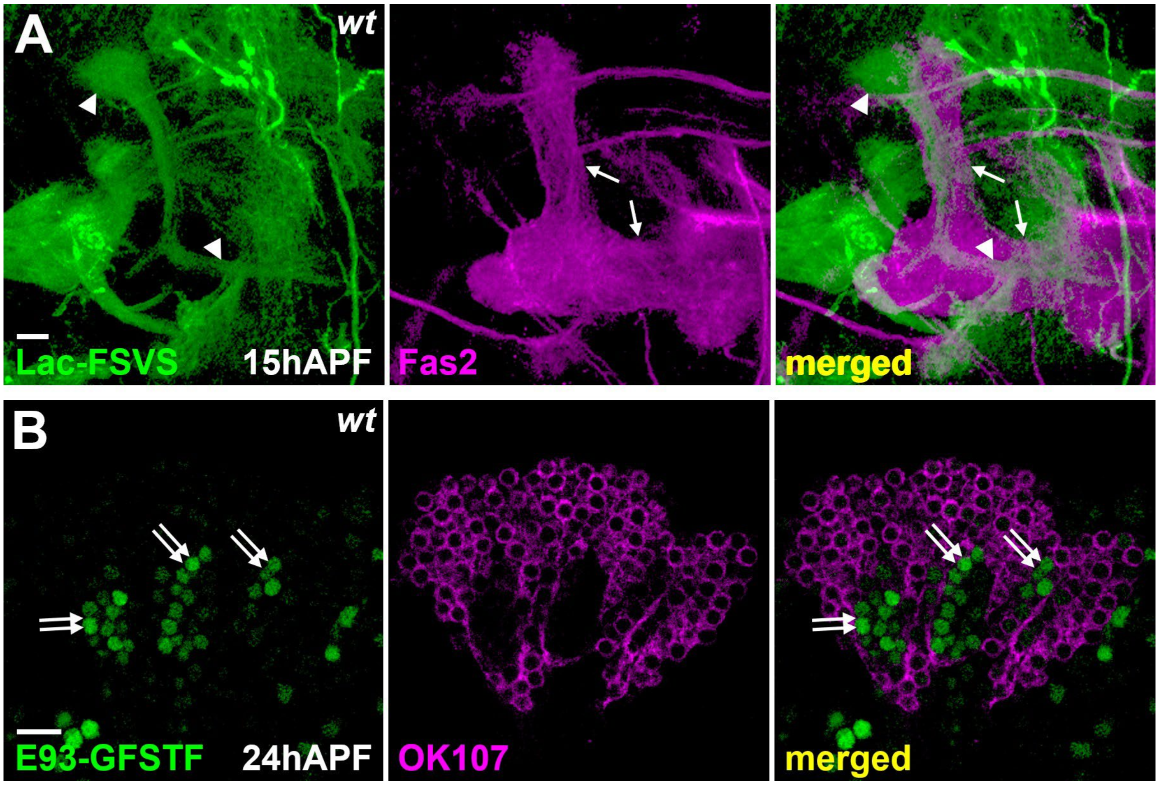


**S3 Fig. Early pupal expression of Lac-FSVS and E93-GFSTF in KCs**

(A-B) The expression of Lac-FSVS (green in panel A) and E93-GFSTF (green in panel B) was observed at 15h and 24h after puparium formation (APF), respectively. Lac-FSVS was expressed in α'/β' neurons (arrowheads) according to counter-staining with cell adhesion molecule Fasciclin II (Fas2, magenta in panel A), which primarily labels γ neurons (arrows) at 15 h APF. E93-GFSTF (double-arrows) was seen in the region with the weak RFP expression driven by GAL4-OK107 (magenta in panel B). This pattern implies that F93-GFSTF expression occurs in the newly generated KCs, which are most likely α/β neurons, at 24 h APF. Scale bar: 10 μm.

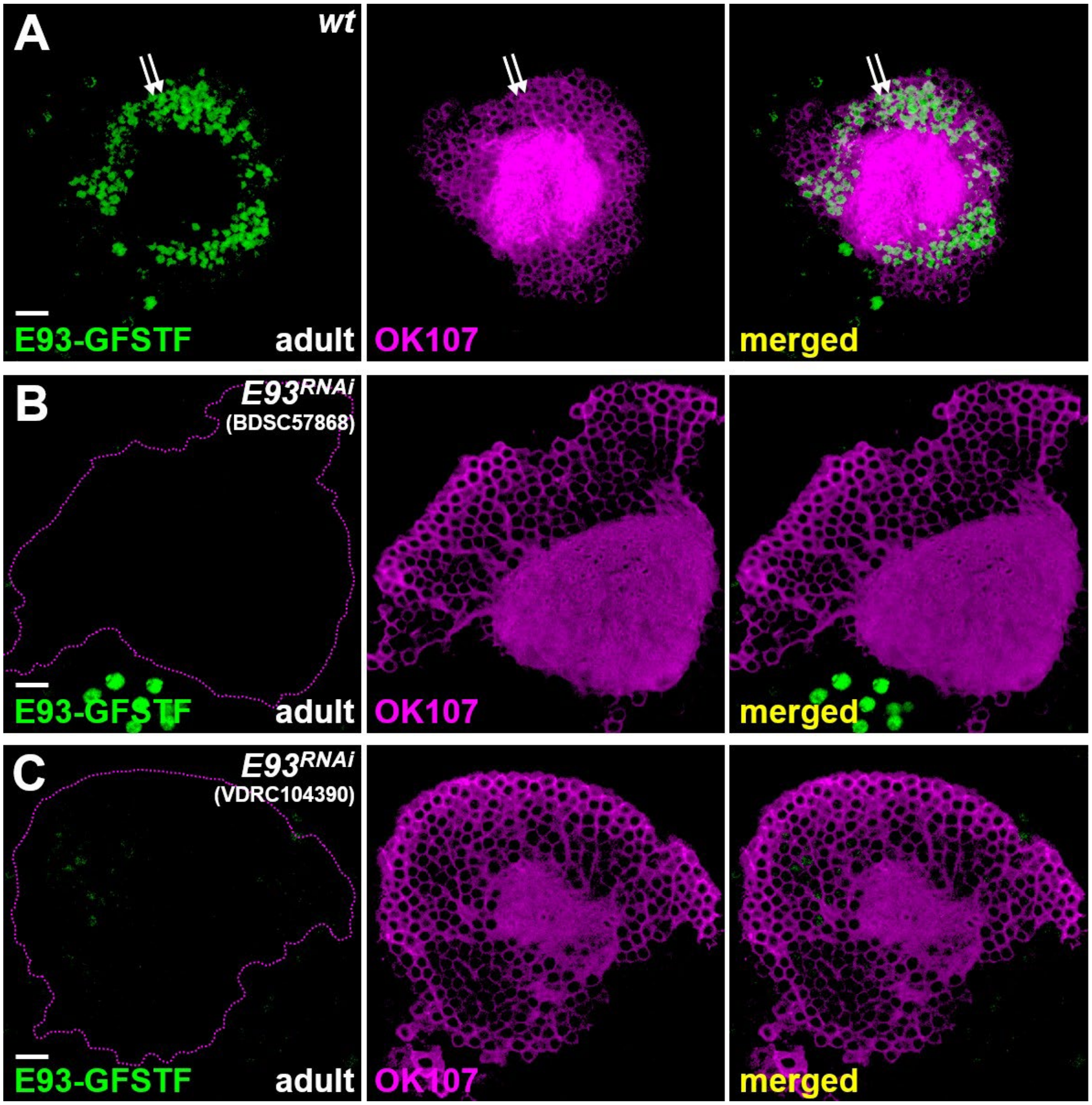


**S4 Fig. Specificity of *E93* RNAi reagents in blocking the expression of E93-GFSTF**

(A-C) Overexpression of either *E93* RNAi [from Bloomington *Drosophila* Stock Center (BDSC) stock number 57868 or Vienna *Drosophila* Resource Center (VDRC) stock number 104390] using GAL4-OK107 (magenta) led to specific knockdown of E93-GFSTF (green) in KCs (double-arrows) in adult brains. *E93* RNAi (BDSC57868) was used in the rest of this study. Scale bar: 10 μm.

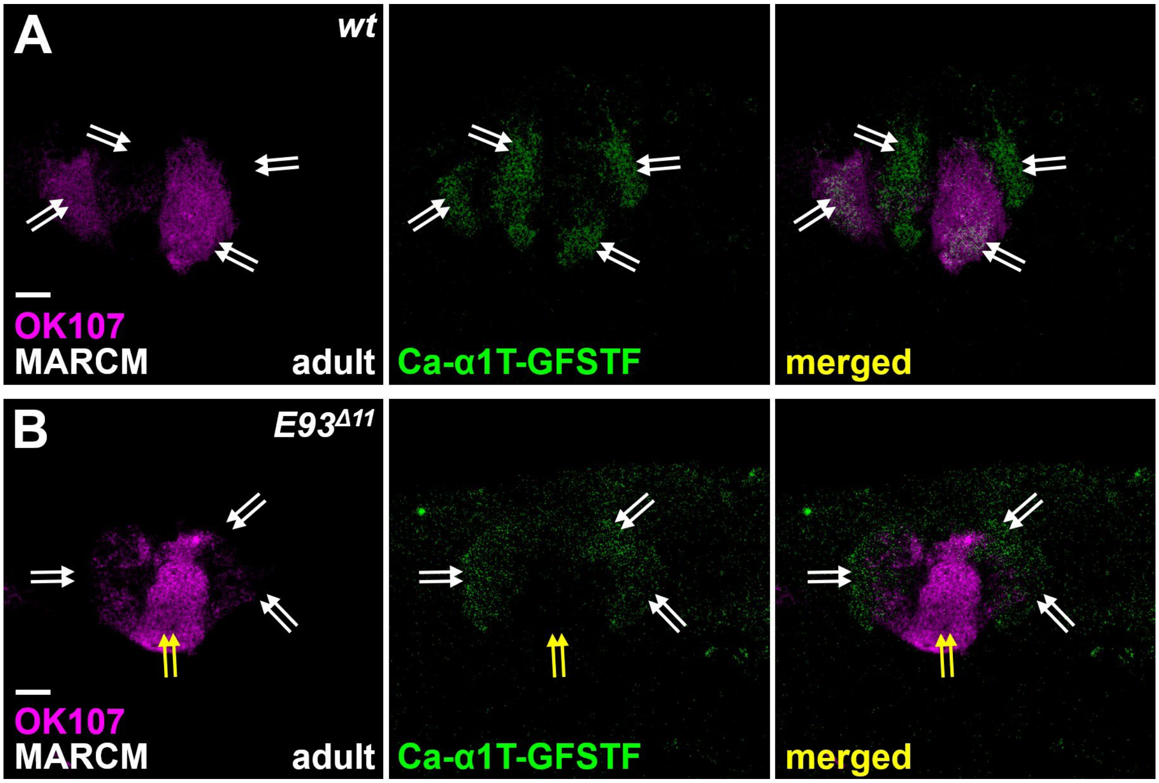


**S5 Fig. Downregulation of Ca-α1T-GFSTF in the *E93* mutation**

(A-B) As compared to the wild-type sample, the Ca-α1T-GFSTF expression (green, indicated by double-arrows) was abolished in the calyx region of $E93^{\Delta 11}$ mutants in the MARCM analysis using GAL4-OK107 (magenta). Mosaic clones were induced in newly hatched larva and analyzed in adult brains. Scale bar: 10 µm.

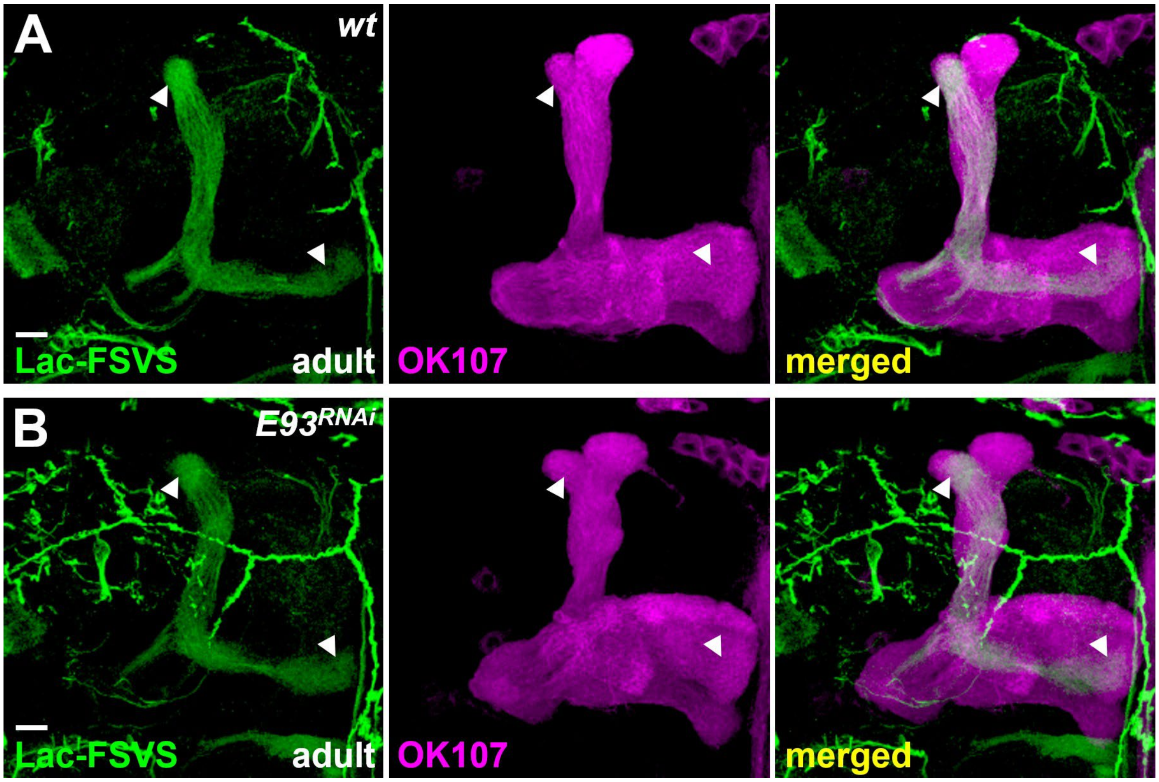


**S6 Fig. *E93* RNAi knockdown does not affect the expression of Lac-FSVS**

Compared to wild-type samples, overexpression of *E93* RNAi driven by GAL4-OK107 (magenta) did not block the expression of Lac-FSVS (green) in α'/β' neurons (arrowheads) in adult brains. Scale bar: 10 μm.

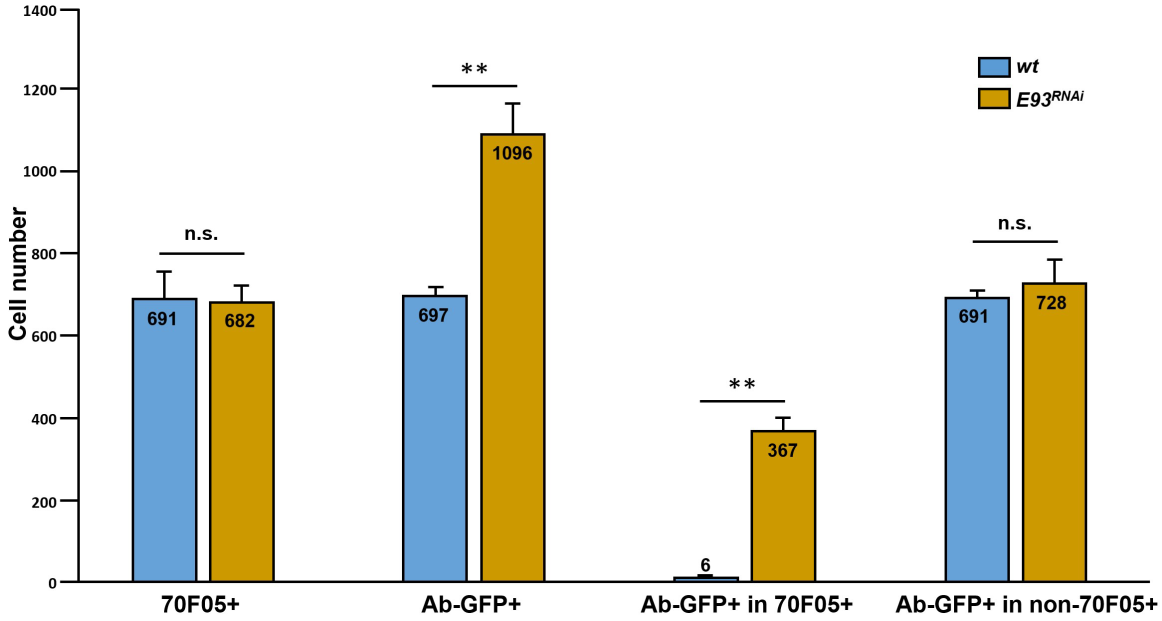


**S7 Fig. Statistical analysis of Ab-GPF expression in KCs of wild-type and *E93* RNAi knockdown samples in Fig 2H and 2I**

Compared to the wild-type (691±67, n=5), *E93* RNAi knockdown driven by GAL4-OK107 did not alter the cell number of 70F05-LexA-positive neurons (682±41, n=6). However, compared to the wild-type (697±24), *E93* RNAi knockdown significantly increased the cell number of Ab-GFP-positive neurons (1096±77). Interestingly, Ab-GFP was rarely expressed in 70F05-LexA-positive neurons in wild-type samples (6±2). In contrast, Ab-GFP was expressed in more than half of all 70F05-LexA-positive neurons in *E93* RNAi knockdown samples (367±36). The cell numbers of Ab-GFP outside 70F05-LexA-positive neurons were similar between wild-type (691±24) and *E93* RNAi knockdown (728±60) samples. Student's test was used for statistical analysis. n.s.: not significant; **: significant.

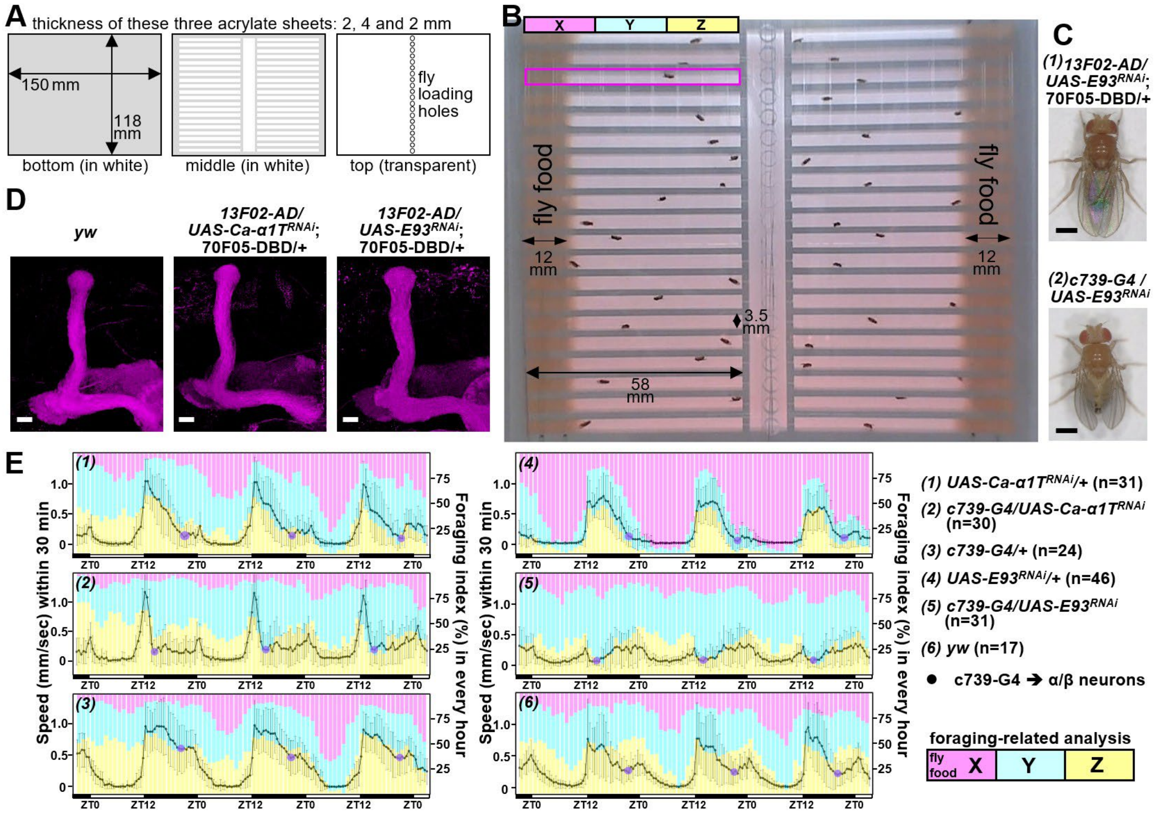


**S8 Fig. Setting, tracking and related results for the behavioral assay**

(A-B) The activity monitor system was developed by DroBot, Inc. The setting of the behavioral assay was assembled by three acrylate sheets (sizes and properties as depicted). Flies were loaded using the holes on the top plate to set up left and right experimental groups. Before loading flies, food was pre-placed on distal sides of chambers. Individual fly activities were filmed for more than four days. Images derived from videos were analyzed by the built-in software of the DroBot activity monitor system. Average speed and standard deviation of individual flies were calculated according to total traveling distance in 30 min. (C) Samples of *E93* knockdown driven by c739-GAL4 displayed curly and aberrant wings, which might compromise the general movement seen in panel E(5). (D) MB lobes appeared intact when *Ca-α1T* and *E93* were knocked down by an α/β neural driver, 13F02-AD/70F05-DBD. (E) The overall pattern of moving speed in RNAi knockdown samples of *Ca-α1T* and *E93* driven by GAL4-c739 (another α/β neural driver) was similar to those samples used 13F02-AD/70F05-DBD (seen in Fig 2I). Overall moving speed was lower and more variable in *E93* knockdowns driven by GAL4-c739, possibly resulting from the aberrant wings seen in panel C(2). Compared to control flies, *Ca-α1T* and *E93* knockdown flies (with GAL4-c739) explored less in the X zone, especially on day four. ZT: Zeitgeber time. Scale bar: 0.5 mm in panel C and 10 μm in panel D.

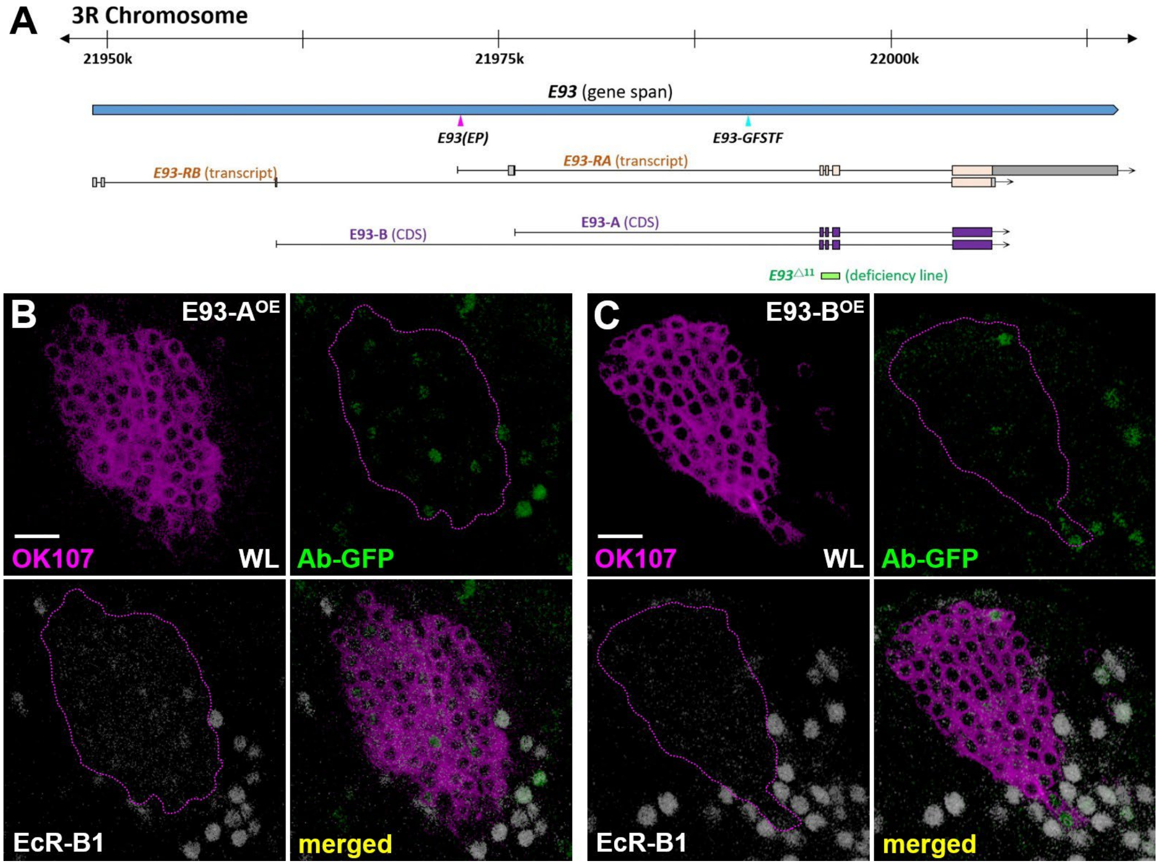


**S9 Fig. *E93* gene and downregulation of Ab-GFP and EcR-B1 in KCs by overexpression of E93 isoforms**

(A) Based on the information available at Flybase, the *E93* gene potentially expresses two *E93* transcript variants that encode E93-A and E93-B protein isoforms. E93-A and E93-B isoforms share most of the protein sequence but differ in respective 9 and 32 unique amino acids at the N-terminus. The E93(EP) is inserted at the proximal region of the *E93-A* 5'UTR, was used to overexpress E93 in most of the (GOF) experiments in this paper. The $E93^{\Delta 11}$ mutation is a small deficiency line generated by deleting the genomic DNA from exon 2 to exon 4 [8]. (B-C) The expression of two γ neural-specific genes, Ab-GFP isoforms (green) and EcR-B1 (white), was almost absent in KCs at the WL stage when E93-A and E93-B were overexpressed driven by GAL4-OK107 (magenta). Since similar results were found when overexpressing E93(EP), E93-A and E93-B (Fig 3F), only the E93(EP) line was used for most of gain-of-function studies in this paper. Scale bars: 10 μm.

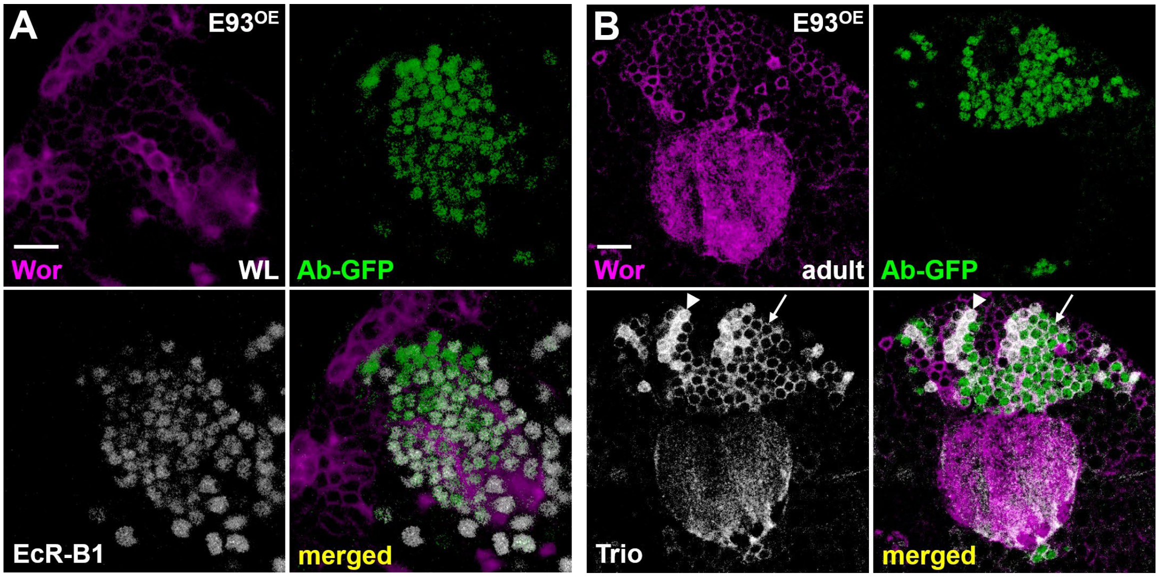


**S10 Fig. E93 overexpression using a neuroblast driver does not cause defects in KCs**

(A-B) The expression of γ neural-specific genes, Ab-GFP isoforms (green), EcR-B1 (white) and cytosolic expressed Trio (white, arrows), was intact in KCs at WL and adult stages when E93(EP) overexpression was driven by a pan-neuroblast driver, Worniu (Wor)-GAL4 (magenta) [9]. Similarly, the whole-cell expression level of Trio, an α'/β' neural-specific maker (white, arrowheads), was not altered at the adult stage by E93 overexpression. These results suggest that defects in KCs caused by E93 overexpression in Fig 3 and S9 Fig are not due to impairments in MB neuroblasts. Scale bars: 10 μm.

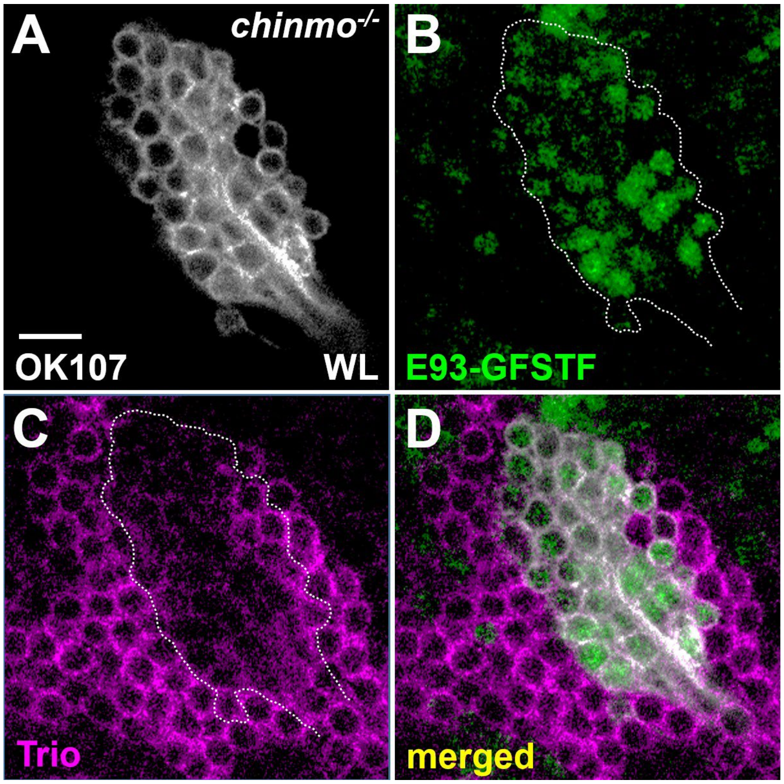


**S11 Fig. Precocious upregulation of E93-GFSTF in KCs in the *chinmo* mutation**

E93-GFSTF expression (green) was precociously turned on in KCs of *chinmo*[1] mutants in the MARCM analysis using GAL4-OK107 (white). Mosaic clones were induced in newly hatched larva and analyzed at the WL stage. The expression of Trio (magenta), labeling γ neurons at the WL stage, was significantly reduced in the *chinmo* mutant clone. Scale bar: 10 μm.

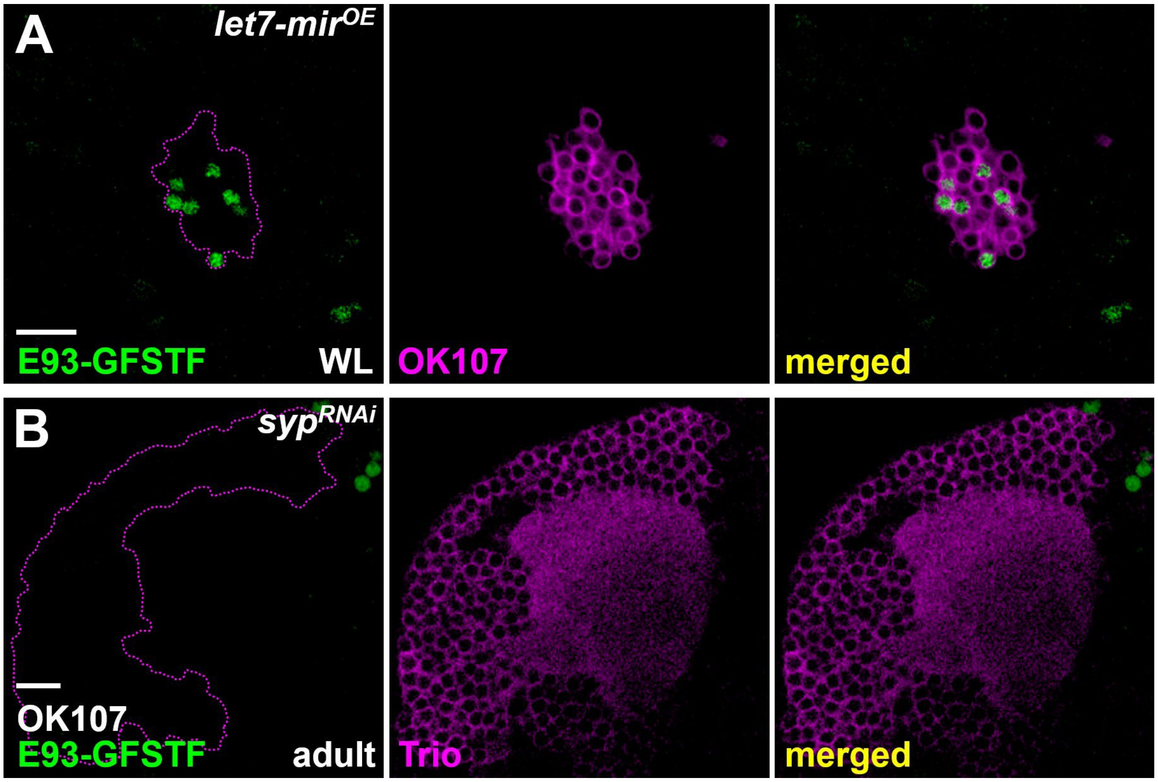


**S12 Fig. Overexpression of *let-7* and *syp* RNAi compromises the expression of E93-GFSTF in KCs**

Overexpression of microRNA *let-7* and RNAi of RNA binding protein *syp* driven by GAL4-OK107 (magenta) led to respective upregulation and downregulation of E93-GFSTF (green) expression in KCs at WL and adult stages, respectively. Scale bar: 10 µm.

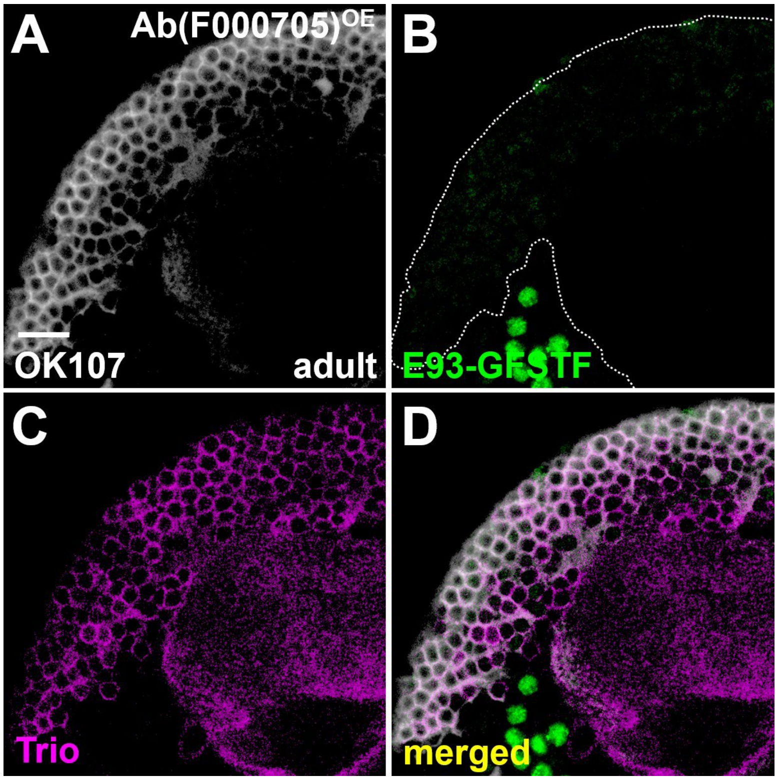


**S13 Fig. Overexpression of Ab compromises E93-GFSTP expression in KCs**

(a-d) Overexpression of Ab (an independent transgenic line from the FlyORF stock center; stock number F000705) driven by GAL4-OK107 (white) significantly blocked E93-GFSTF expression (green) in KCs of adult brains. Similarly, as seen in Fig 4G-H, Trio (magenta) was also expressed in the cytosol in almost all KCs in FlyORF Ab gain-of-function studies. Scale bars: 10 μm.

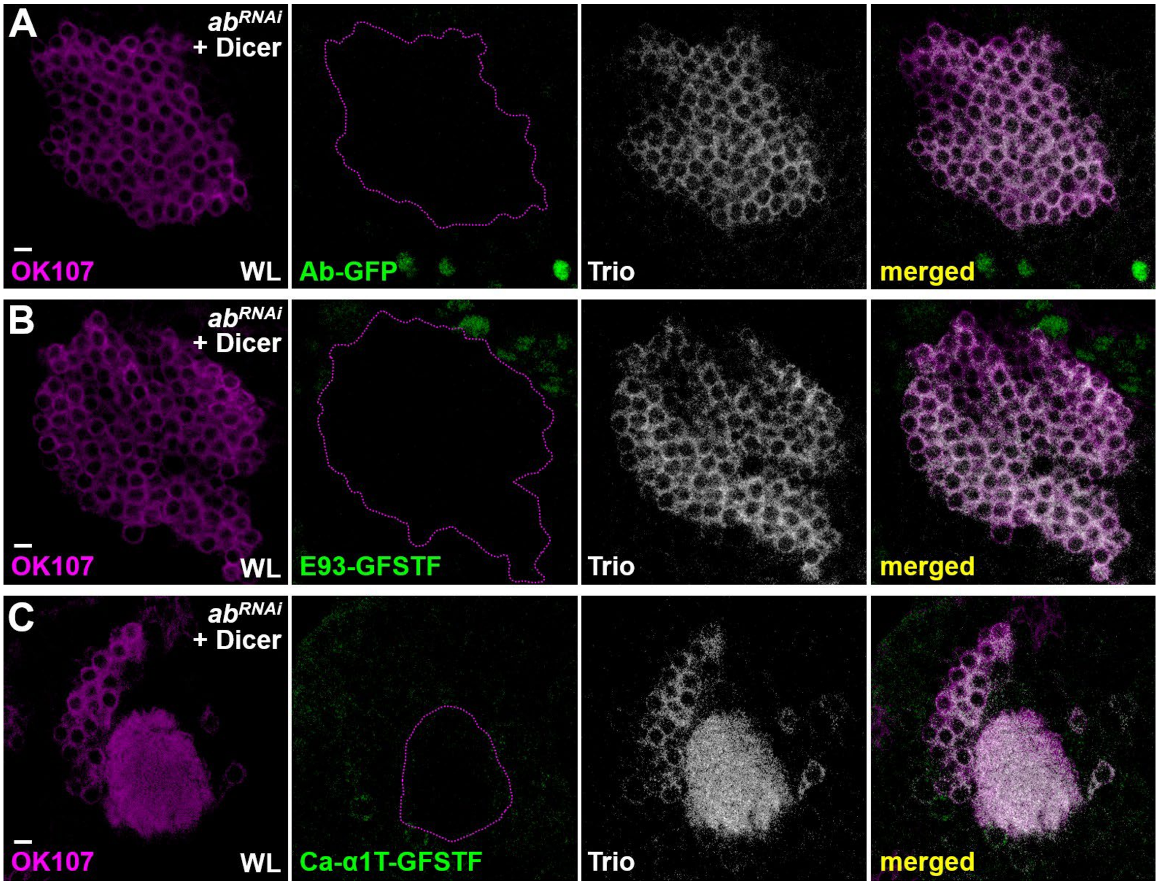


**S14 Fig. RNAi knockdown of *ab* elicits no obvious effects on the expression of Trio, E93-GFSTF and Ca-α1T-GFSTF in KCs**

(A-C) RNAi knockdown of *ab* (available at Vienna *Drosophila* stock center, stock number 104582) using GAL4-OK107 (magenta) specifically blocked the expression of Ab-GFP (green) in KCs at the WL stage. However, *ab* RNAi knockdown neither abolished Trio expression (white) nor did it upregulate expression of E93-GFSTF (green) and Ca-α1T-GFSTF (green) in KCs at the WL stage. Dicer was included to increase the *ab* RNAi knockdown potency in experiments shown in panels A-C. Scale bars: 10 μm.

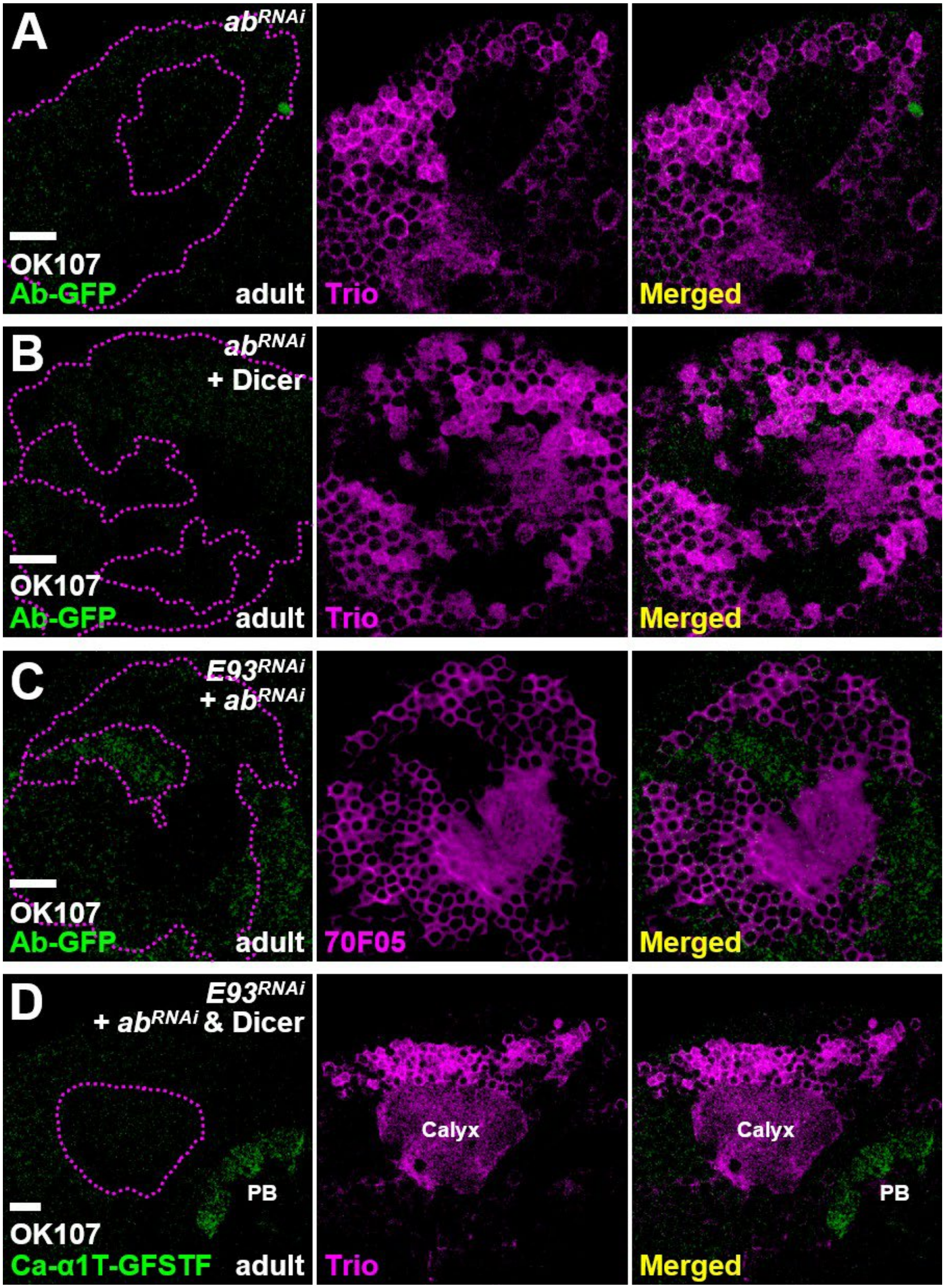


**S15 Fig. RNAi knockdown of *ab* fails to restore the Ca-α1T-GFSTF expression caused by E93 knockdown**

(A-B) RNAi knockdown of *ab* using GAL4-OK107 blocked the expression of Ab-GFP (green) in KCs of adult brains. (C-D) Ca-α1T-GFSTF expression (green) was not restored in KCs with double RNAi knockdown of *E93* and *ab* at the adult stage, while Ab-GFP upregulation caused by *E93* knockdown was abolished by overexpressing *ab* RNAi. Dicer was included to increase the *ab* RNAi knockdown potency in experiments shown in panels B and D. Trio expression was unaltered in experiments shown in panels A, B and D. 70F05-LexA-posive cells were used to test whether *ab* RNAi could block the upregulation of Ab-GFP expression caused by *E93* knockdown in panel c. Scale bars: 10 μm.

### Materials and Methods

#### Experimental model and subject details

Flies were cultured in a room maintained at 25ºC (± 1.5ºC) and 50-65% humidity for all experiments. For most experiments, flies were used with no selection for sex; therefore, roughly equal numbers of males and females were used. However, females were selected for the analysis of Ca-α1T-GFSTF in Fig. 1G-H, 2A-B, 3A-B and 4I-J and S5A-B, S13 and S14D Figs due to the cytolocation of the *Ca-α1T* gene on the X chromosome. In addition, males were selected for the behavioral assay to mitigate complications of egg laying behaviors on locomotion tracking.

#### Fly strains

The fly strains used in this study were as follows. Most strains are available from either Bloomington *Drosophila* stock center (BDSC), Kyoto *Drosophila* stock center (DGGR) or Vienna *Drosophila* stock center (VDRC). (1) *Ab-GFP* (BDSC38626); (2) *Lac-FSVS* (DGGR115308); (3) *E93-GFSTF* (BDSC59412); (4) *Ca-α1T-GFSTF* (BDSC61800); (5) *hs-FLP*$^{12}$*,UAS-mCD8::GFP* (BDSC28832); (6) *tubP-GAL80,FRT*$^{40A}$ (BDSC5192); (7) *chinmo*$^{[1]}$*,FRT*$^{40A}$ (BDSC59969); (8) *GAL4-OK107* (BDSC854); (9) *UAS-mCD8::RFP [attp40]* (BDSC32219); (10) *UAS-mCD8::RFP [attp2]* (BDSC32218); (11) *UAS-E93 RNAi* (BDSC57868); (12) *UAS-E93 RNAi* (VDRC104390); (13) *UAS-Ca-α1T RNAi* (BDSC39029); (14) *GAL4-c739* (BDSC7362); (15) *44E04-LexA::P65* (BDSC52736); (16) *70F05-LexA::P65* (BDSC523629); (17) *lexAop2-myr::GFP* [10]; (18) *lexAop2-mCD8::RFP* [10]; (19) *FRT*$^{82B}$*,tubP-GAL80* (BDSC5135); (20) *FRT*$^{82B}$ (BDSC86313); (21) *E93*$^{\Delta 11}$ (BDSC93128); (22) *E93(EP)* (BDSC30179); (23) *UAS-E93-A [VK37]* (this study); (24)

*UAS-E93-B [VK37]* (this study); (25) *GAL4-201Y* (BDSC4440); (26) *mamo*$^{H/I}$*-HA*[11]; (27) *mamo*$^{D\sim G}$*-HA* [11]; (28) *UAS-chinmo RNAi [VK37]* [4]; (29) *UAS-LUC-let7* (BDSC41171); (30) *UAS-Ssyp RNAi* (VDRC33011); (31) *UAS-mamo RNAi* (BDSC44103); (32) *UAS-ab* (BDSC23639); (33) *UAS-ab-HA* (FlyORF000705); (34) *UAS-Dcr2* (BDSC24651); (35) *UAS-ab RNAi* (VDRC104582); (36) *13F02-p65.AD,44E04-GAL4.DBD* (BDSC68291); (37) *70F05-GAL4.DBD* (BDSC69380). The *UAS-E93-A* and *UAS-E93-B* transgenes were generated using standard molecular biology methods to clone cDNA fragments derived from fully-sequenced EST clones, GH10557 and LP08695 (available from *Drosophila* Genomics Resource Center), carrying *E93-A* and *E93-B* isoforms into the attB-UAST vector. The generation of *UAS-E93-A* and *UAS-E93-B* transgenes and fly stains was performed by WellGenetics, Inc.

**RNAi knockdown and overexpression experiments and MARCM clonal analyses**

*UAS-RNAi* and *UAS-transgene* lines were crossed to GAL4-107 and GAL4-201Y for knockdown and overexpression of genes of interest in KCs. Mosaic clones for the MARCM studies were generated as previously described [12]. In short, mosaic clones of *chinmo*$^{[1]}$ and *E93*$^{\Delta 11}$ mutations were induced by 35 min of heat shock using *hs-FLP*$^{[12]}$ in newly hatched larva. Dissection, immunostaining and mounting of adult brains were performed as described in a standard protocol [12]. Primary antibodies used in this study included guinea pig antibody against Chinmo (1:1000, Sokol laboratory [13]), rat monoclonal antibody against mCD8 (1:100, Thermo Fisher Scientific), rabbit antibody against GFP (1:750, Thermo Fisher Scientific), and mouse monoclonal antibodies against EcR-B1 (1:50, DSHB), Fas2 (1:100, DSHB) and Trio (1:50, DSHB). Secondary antibodies conjugated to different fluorophores (Alexa 488, Alexa 546 and Alexa 647; Thermo Fisher Scientific and Jackson ImmunoResearch Lab, Inc.) were used at 1:750 dilutions. Immunofluorescence images were collected by confocal microscopy on a Zeiss LSM 700, projected using the LSM browser and processed in Adobe Photoshop CS6. All data are representative of more than 3 brains per genotype.

**Distance measurement in the behavioral assay and statistical analysis**

Individual fly activities were recorded and analyzed using the activity monitor system developed by DroBot, Inc. This system used an open-resourced software pySolo to track individual moving flies [14]. Average speed and standard deviation of individual flies were calculated according to total traveling distance within 30 min periods from the first day to the fifth day, as shown in Fig 2I and S8E Fig. Student's t-test was used for statistical analysis to compare datasets with two groups in S8 Fig.

### Supplementary Table 1. Expression patterns of GFP lines in S1 Fig

| panel | gene | stock # | in KCs at WL | in KCs at adult |
|---|---|---|---|---|
| A1 | *dan* | BDSC92324 | no | no |
| A2 | *dlp* | BDSC60540 | calyx and lobe of KCs | enriched in calyx and lobe of γ |
| A3 | *ed* | BDSC59777 | no | no |
| A4 | *Imp* | DGGR115455 | no | cytosol of γ and α'/β' |
| A5 | *SIFaR* | BDSC60228 | no | no |
| A6 | *TkR86C* | BDSC60549 | no | no |
| B1 | *CG31637* | BDSC64438 | no | no |
| B2 | *CG43373* | BDSC60239 | no | no |
| B3 | *CG4404* | BDSC90835 | no | no |
| B4 | *crb* | BDSC61781 | no | no |
| B5 | *Lmpt* | BDSC66776 | no | no |
| B6 | *mbc* | DGGR115505 | no | no |
| B7 | *Octβ3R* | BDSC60245 | no | no |
| C1 | *Ace* | BDSC60260 | calyx of KCs | calyx of KCs |
| C2 | *app* | BDSC60283 | no | no |
| C3 | *beat-IV* | BDSC66506 | KCs | γ and α'/β' |
| C4 | *Ccn* | BDSC60259 | no | no |
| C5 | *CG4829* | DGGR115623 | no | enriched in α'/β' |
| C6 | *Cyp4p3* | BDSC59829 | no | cytosol and calyx of KCs |
| C7 | *DAT* | VDRC318840 | MB lobe (maybe not KCs) | MB lobe (maybe not KCs) |
| C8 | *dnr1* | BDSC76236 | no | no |
| C9 | *dpr17* | BDSC61801 | no | no |
| C10 | *Epac* | BDSC66364 | no | no |
| C11 | *eys* | BDSC63162 | no | no |
| C12 | *fz3* | VDRC318166 | no | no |
| C13 | *igl* | BDSC60527 | KCs | KCs |
| C14 | *LRP1* | BDSC60248 | no | no |
| C15 | *mamo* | VDRC318601 | no | no |
| C16 | *Mp* | BDSC60567 | no | cytosol and calyx of KCs |
| C17 | *msi* | BDSC61750 | no | enriched in cell body of α'/β' |
| C18 | *Ndae1* | BDSC61778 | no | no |
| C19 | *nuf* | BDSC61802 | no | no |
| C20 | *rhea* | BDSC39649 | no | no |
| C21 | *smal* | VDRC318203 | no | no |
| C22 | *tok* | BDSC60550 | no | no |
| C23 | *Zasp67* | VDRC318355 | no | no |

### Supplementary Table 2. Genotypes of flies shown in each figure panel

| Figure | Genotype |
|---|---|
| 1A,1B | *w;+;ab-GFP+;+* |
| 1C,1D,S3A | *w;Lac-FSVS/+;+;+* |
| 1E,1F | *yw;+;E93-GFSTF/+;+* |
| 1G,1H | *yw,Ca-α1T-GFSTF/+;+;+;+* |
| 2A,3A,4I | *yw,Ca-α1T-GFSTF/w;+;UAS-mCD8::RFP/+;GAL4-OK107/+* |
| 2B | *yw,Ca-α1T-GFSTF/w;UAS-E93-RNAi*$^{BDSC57868}$*/+;UAS-mCD8::RFP/+;GAL4-OK107/+* |
| 2C | *w;44E04-LexA::P65/+;UAS-mCD8::RFP,lexAop-myr::GFP/+;GAL4-OK107/+* |
| 2D | *w;44E04-LexA::P65/UAS-E93-RNAi*$^{BDSC57868}$*;UAS-mCD8::RFP,lexAop-myr::GFP/+;GAL4-OK107/+* |
| 2E | *w;70F05-LexA::P65/+;UAS-mCD8::RFP,lexAop-myr::GFP/+;GAL4-OK107/+* |
| 2F | *w;70F05-LexA::P65/UAS-E93-RNAi*$^{BDSC57868}$*;UAS-mCD8::RFP,lexAop-myr::GFP/+;GAL4-OK107/+* |
| 2G | *w;70F05-LexA::P65/+;lexAop-mCD8::RFP/Ab-GFP;GAL4-OK107/+* |
| 2H | *w;70F05-LexA::P65/UAS-E93-RNAi*$^{BDSC57868}$*;lexAop-mCD8::RFP/Ab-GFP;GAL4-OK107/+* |
| 2I(1),S8E(1) | *yw,UAS-Ca-α1T RNAi*$^{BDSC39029}$*/+;+;+;+* |
| 2I(2), S8D(2) | *yw,13F02-AD/UAS-Ca-α1T RNAi*$^{BDSC39029}$*;70F05-DBD/+;+* |
| 2I(3) | *yw,13F02-AD/+;70F05-DBD/+;+* |
| 2I(4),S8E(4) | *yw,UAS-E93-RNAi*$^{BDSC57868}$*/+;+;+;+* |
| 2I(5),S8C(1),S8D(3) | *yw,13F02-AD/UAS-E93-RNAi*$^{BDSC57868}$*;70F05-DBD/+;+* |
| 2I(6),S8D(1),S8E(6) | *yw,+;+;+* |
| 3B | *yw,Ca-α1T-GFSTF/w;+;UAS-mCD8::RFP/E93(EP);GAL4-OK107/+* |

| | |
|---|---|
| 3C | w;GAL4-201Y,70F05-LexA::P65/+;UAS-mCD8::RFP,lexAop-myr::GFP/+;+ |
| 3D | w;GAL4-201Y,70F05-LexA::P65/+;UAS-mCD8::RFP,lexAop-myr::GFP/E93(EP);+ |
| 3E | w;UAS-mCD8::RFP/+;ab-GFP/+;GAL4-OK107/+ |
| 3F | w;UAS-mCD8::RFP/+;ab-GFP/E93(EP);GAL4-OK107/+ |
| 3G | w,mamo$^{H/I}$-HA/w;+;UAS-mCD8::GFP/+;GAL4-OK107/+ |
| 3H | w,mamo$^{H/I}$-HA/w;+;UAS-mCD8::GFP/E93(EP);GAL4-OK107/+ |
| 3I | w,mamo$^{D\sim G}$-HA/w;+;UAS-mCD8::GFP/+;GAL4-OK107/+ |
| 3J | w,mamo$^{D\sim G}$-HA/w;+;UAS-mCD8::GFP/E93(EP);GAL4-OK107/+ |
| 3K,S6A | w;Lac-FSVS/+;UAS-mCD8::RFP/+;GAL4-OK107/+ |
| 3L | w;Lac-SVS/+;UAS-mCD8::RFP/E93(EP);GAL4-OK107/+ |
| 4A | w;+;UAS-mCD8::RFP/Ab-GFP;GAL4-OK107/+ |
| 4B | w;UAS-E93-RNAi$^{BDSC57868}$/+;UAS-mCD8::RFP/Ab-GFP;GAL4-OK107/+ |
| 4C,4F,S3B, S4A | yw/w;UAS-mCD8::RFP/+;E93-GFSTF/+;GAL4-OK107/+ |
| 4D | yw/w;UAS-mCD8::RFP/UAS-chinmo-RNAi;E93-GFSTF/+;GAL4-OK107/+ |
| 4E,4G | yw/w;UAS-mCD8::RFP/UAS-mamo-RNAi;E93-GFSTF/+;GAL4-OK107/+ |
| 4H | yw/w;UAS-mCD8::RFP/+;E93-GFSTF/UAS-ab $^{BDSC23639}$;GAL4-OK107/+ |
| 4J | yw,Ca-α1T-GFSTF/w;+;UAS-mCD8::RFP/UAS-ab $^{BDSC23639}$;GAL4-OK107/+ |
| S1 | The genotype can be found by stock number listed in Supplementary Table 2 |
| S2 | hs-FLP$^{[122]}$/w;chinmo$^{1}$,FRT$^{40A}$/tubP-GAL80,FRT$^{40A}$;Ab-GFP/UAS-mCD8::RFP;GAL4-OK107/+ |
| S4B | yw/w;UAS-mCD8::RFP/UAS-E93-RNAi$^{BDSC57868}$;E93-GFSTF/+;GAL4-OK107/+ |
| S4C | yw/w;UAS-mCD8::RFP/UAS-E93-RNAi$^{VDRC104390}$;E93-GFSTF/+;GAL4-OK107/+ |
| S5A | yw,Ca-α1T-GFSTF/hs-FLP$^{[122]}$;UAS-mCD8::RFP/+;FRT$^{82B}$/FRT$^{82B}$,tubP-GAL80;GAL4-OK107/+ |
| S5B | yw,Ca-α1T-GFSTF/hs-FLP$^{[122]}$;UAS-mCD8::RFP/+;FRT$^{82B}$,E93$^{\Delta 11}$/FRT$^{82B}$,tubP-GAL80;GAL4-OK107/+ |

| | |
|---|---|
| S6B | w;Lac-SVS/UAS-E93-RNAi$^{BDSC57868}$;UAS-mCD8::RFP/+;GAL4-OK107/+ |
| S8E(2) | yw,GAL4-c739/UAS-Ca-α1T RNAi$^{BDSC39029}$;+;+ |
| S8E(3) | yw,GAL4-c739/+;+;+ |
| S8C(2),<br>S8E(5) | yw,GAL4-c739/UAS-E93-RNAi$^{BDSC57868}$;+;+ |
| S9B | w;UAS-mCD8::RFP/UAS-E93-A;ab-GFP/+;GAL4-OK107/+ |
| S9C | w;UAS-mCD8::RFP/UAS-E93-B;ab-GFP/+;GAL4-OK107/+ |
| S10A,S10B | w;wor-GAL4,UAS-mCD8::RFP/+;ab-GFP/E93(EP);+ |
| S11 | hs-FLP$^{[122]}$/w;chinmo$^{1}$,FRT$^{40A}$/tubP-GAL80,FRT$^{40A}$;E93-GFSTF/UAS-mCD8::RFP;GAL4-OK107/+ |
| S12A | w;UAS-mCD8::RFP/+;E93-GFSTF/UAS-LUC-let7;GAL4-OK107/+ |
| S12B | w;UAS-mCD8::RFP/UAS-syp-RNAi$^{VDRC33011}$;E93-GFSTF/+;GAL4-OK107/+ |
| S13 | yw/w;UAS-mCD8::RFP/+;E93-GFSTF/UAS-ab$^{F000705}$;GAL4-OK107/+ |
| S14A,S15B | w;UAS-mCD8::RFP/UAS-ab-RNAi$^{VDRC104582}$;ab-GFP/UAS-Dcr2.0;GAL4-OK107/+ |
| S14B | w;UAS-mCD8::RFP/UAS-ab-RNAi$^{VDRC104582}$;E93-GFSTF/UAS-Dcr2.0;GAL4-OK107/+ |
| S14C | yw,Ca-α1T-GFSTF/w;UAS-mCD8::RFP/UAS-ab-RNAi$^{VDRC104582}$;UAS-Dcr2.0/+;GAL4-OK107/+ |
| S15A | w;UAS-mCD8::RFP/UAS-ab-RNAi$^{VDRC104582}$;ab-GFP/+;GAL4-OK107/+ |
| S15C | w;70F05-LexA/UAS-E93-RNAi$^{BDSC57868}$,UAS-ab-RNAi$^{VDRC104582}$;lexAop-mCD8::RFP/ab-GFP;GAL4-OK107/+ |
| S15D | yw,Ca-α1T-GFSTF/UAS-E93-RNAi$^{BDSC57868}$,UAS-ab-RNAi$^{VDRC104582}$;UAS-mCD8::RFP/UAS-Dcr2.0;GAL4-OK107/+ |

### Key resources table

| REAGENT or RESOURCE | SOURCE | IDENTIFIER |
|---|---|---|
| **Antibodies** | | |
| Mouse anti-Fas2 | DSHB | Cat# AB_528235; RRID: AB_528235 |
| Mouse anti-EcR-B1 | DSHB | Cat# AB_2154902; RRID: AB_2154902 |
| Mouse anti-Trio | DSHB | Cat# AB_528494; RRID: AB_528494 |
| Rat anti-CD8 | Thermo Fisher Scientific | Cat# MCD0800; RRID: AB_10392843 |
| Rat anti-HA | Roche | Cat# 11867423001; RRID: AB_390918 |
| Rabbit anti-GFP | Thermo Fisher Scientific | Cat# A-11122; RRID: AB_221569 |
| Guinea pig anti-Chinmo | Sokol Lab | 13 |
| Goat anti-rabbit Alexa 488 | Thermo Fisher Scientific | Cat# A-11034; RRID: AB_2576217 |
| Goat anti-rabbit Alexa 546 | Thermo Fisher Scientific | Cat# A-11081; RRID: AB_2534125 |
| Goat anti-guinea pig Alexa 647 | Thermo Fisher Scientific | Cat# A-21450; RRID: AB_2535867 |
| Goat anti-mouse Alexa 647 | Jackson ImmunoResearch lab, Inc. | Cat# 115-605-166; RRID: AB_2338914 |
| **Chemicals, peptides and recombinant proteins** | | |
| Formaldehyde 37% solution | Sigma-Aldrich | Cat# 252549 |
| Paraformaldehyde 16% solution | Electron Microscopy Sciences | Cat# 15710 |
| SlowFade$^{TM}$ Gold Antifade Mountant | Thermo Fisher Scientific | Cat# S36936 |
| **Experimental models: Organisms/strains** | | |
| *D. melanogaster*: *Ab-GFP [VK33]* | Bloomington *Drosophila* Stock Center (BDSC) | BDSC_38626 |
| *D. melanogaster*: *Lac-FSVS* | Kyoto *Drosophila* Resource Center (DGGR) | DGGR_115308 |
| *D. melanogaster*: *E93-GFSTF* | BDSC | BDSC_59412 |
| *D. melanogaster*: *Ca-α1T-GFSTF* | BDSC | BDSC_61800 |
| *D. melanogaster*: *hs-FLP[12],UAS-mCD8::GFP* | BDSC | BDSC_28832 |
| *D. melanogaster*: *tubP-GAL80,FRT$^{40A}$* | BDSC | BDSC_5192 |
| *D. melanogaster*: *chinmo$^{1}$,FRT$^{40A}$* | BDSC | BDSC_59969 |
| *D. melanogaster*: *GAL4-OK107* | BDSC | BDSC_854 |
| *D. melanogaster*: *UAS-mCD8::RFP [attP40]* | BDSC | BDSC_32219 |
| *D. melanogaster*: *UAS-mCD8::RFP [attP2]* | BDSC | BDSC_32218 |
| *D. melanogaster*: *UAS-E93-RNAi$^{BDSC57868}$ [attP40]* | BDSC | BDSC_57868 |
| *D. melanogaster*: *UAS-E93-RNAi$^{VDRC104390}$ [VIE-260B]* | Vienna *Drosophila* Resource Center (VDRC) | VDRC_104390 |
| *D. melanogaster*: *UAS-Ca-α1T-RNAi [attP2]* | BDSC | BDSC_39029 |
| *D. melanogaster*: *GAL4-c739* | BDSC | BDSC_7362 |
| *D. melanogaster*: *44E04-LexA::P65 [attP40]* | BDSC | BDSC_52736 |
| *D. melanogaster*: *70F05-LexA::P65 [attP40]* | BDSC | BDSC_53629 |

| | | |
|---|---|---|
| *D. melanogaster*: *13F02-p65.AD [attP40], 44E04-GAL4.DBD [attP2]* | BDSC | BDSC_68291 |
| *D. melanogaster*: *70F05-GAL4.DBD [attP2]* | BDSC | BDSC_69380 |
| *D. melanogaster*: *LexAop2-myr::GFP [VK5]* | Rubin Lab | 10 |
| *D. melanogaster*: *LexAop2-mCD8::RFP [attP2]* | Rubin Lab | 10 |
| *D. melanogaster*: *FRT*$^{82B}$*,tubP-GAL80* | BDSC | BDSC_5135 |
| *D. melanogaster*: *FRT*$^{82B}$ | BDSC | BDSC_86313 |
| *D. melanogaster*: *E93*$^{\Delta 11}$ | BDSC | BDSC_93128 |
| *D. melanogaster*: *E93(EP)* | BDSC | BDSC_30179 |
| *D. melanogaster*: *UAS- E93-A [VK37]* | Yu Lab | this study |
| *D. melanogaster*: *UAS- E93-B [VK37]* | Yu Lab y | this study |
| *D. melanogaster*: *GAL4-201Y* | BDSC | BDSC_4440 |
| *D. melanogaster*: *mamo*$^{H/I}$*-HA* | Yu Lab | 11 |
| *D. melanogaster*: *mamo*$^{D\sim G}$*-HA* | Yu Lab | 11 |
| *D. melanogaster*: *UAS-chinmo-RNAi [VK37]* | Yu Lab | 4 |
| *D. melanogaster*: *UAS-LUC-let7 [attp2]* | BDSC | BDSC_41171 |
| *D. melanogaster*: *UAS-syp-RNAi* | VDRC | VDRC_33011 |
| *D. melanogaster*: *UAS-mamo-RNAi*$^{BDSC44103}$ *[attP40]* | BDSC | BDSC_44103 |
| *D. melanogaster*: *UAS-ab* | BDSC | BDSC_23639 |
| *D. melanogaster*: *UAS-ab-HA [ZH-86Fb]* | Zurich ORFeome Project (FlyORF) | FlyORF_000705 |
| *D. melanogaster*: *UAS-Dcr2* | BDSC | BDSC_24651 |
| *D. melanogaster*: *UAS-ab-RNAi* | VDRC | VDRC_104582 |
| *D. melanogaster*: *dan-GFP [attP40]* | BDSC | BDSC_92324 |
| *D. melanogaster*: *dlp-GFSTF* | BDSC | BDSC_60540 |
| *D. melanogaster*: *ed-GFSTF* | BDSC | BDSC_59777 |
| *D. melanogaster*: *Imp SVS* | DGGR | DGGR_115455 |
| *D. melanogaster*: *SIFaR-GFSTF* | BDSC | BDSC_60228 |
| *D. melanogaster*: *TkR86C-GFSTF* | BDSC | BDSC_60549 |
| *D. melanogaster*: *CG31637-GFSTF* | BDSC | BDSC_64438 |
| *D. melanogaster*: *CG43373-GFSTF* | BDSC | BDSC_60239 |
| *D. melanogaster*: *CG4404-GFP [attP40]* | BDSC | BDSC_90835 |
| *D. melanogaster*: *crb-GFSTF* | BDSC | BDSC_61781 |
| *D. melanogaster*: *Lmpt-GFSTF* | BDSC | BDSC_66776 |
| *D. melanogaster*: *mbc-SVS* | DGGR | DGGR_115505 |
| *D. melanogaster*: *Octbeta3R -GFSTF* | BDSC | BDSC_60245 |
| *D. melanogaster*: *Ace-GFSTF* | BDSC | BDSC_60260 |
| *D. melanogaster*: *app-GFSTF* | BDSC | BDSC_60283 |
| *D. melanogaster*: *beat-IV-GFSTF* | BDSC | BDSC_66506 |
| *D. melanogaster*: *Ccn-GFSTF* | BDSC | BDSC_60259 |
| *D. melanogaster*: *CG4829-FSVS* | DGGR | DGGR_115623 |
| *D. melanogaster*: *Cyp4p3-GFSTF* | BDSC | BDSC_59829 |
| *D. melanogaster*: *DAT-sfGFP [VK33]* | VDRC | VDRC_318840 |
| *D. melanogaster*: *dnr1-GFSTF* | BDSC | BDSC_76236 |
| *D. melanogaster*: *dpr17-GFSTF* | BDSC | BDSC_61801 |
| *D. melanogaster*: *Epac-GFSTF* | BDSC | BDSC_66364 |
| *D. melanogaster*: *eys-GFSTF* | BDSC | BDSC_63162 |
| *D. melanogaster*: *fz3-sfGFP [VK33]* | VDRC | VDRC_318166 |
| *D. melanogaster*: *igl-GFSTF* | BDSC | BDSC_60527 |

| | | |
|---|---|---|
| *D. melanogaster*: *LRP1-GFSTF* | BDSC | BDSC_60248 |
| *D. melanogaster*: *mamo-sfGFP [VK33]* | VDRC | VDRC_318601 |
| *D. melanogaster*: *Mp-GFSTF* | BDSC | BDSC_60567 |
| *D. melanogaster*: *msi-GFSTF* | BDSC | BDSC_61750 |
| *D. melanogaster*: *Ndae1-GFSTF* | BDSC | BDSC_61778 |
| *D. melanogaster*: *nuf-GFSTF* | BDSC | BDSC_61802 |
| *D. melanogaster*: *rhea-GFSTF* | BDSC | BDSC_39649 |
| *D. melanogaster*: *smal-sfGFP [VK33]* | VDRC | VDRC_318203 |
| *D. melanogaster*: *tok-GFSTF* | BDSC | BDSC_60550 |
| *D. melanogaster*: *Zasp67-sfGFP [VK33]* | VDRC | VDRC_318355 |
| **Software and algorithms** | | |
| LSM | Zeiss | N/A |
| Photoshop CS6 | Adobe | N/A |
